# immuneKG: An Immune-Cell-Aware Knowledge Graph Framework for Target Discovery in Immune-Mediated Diseases

**DOI:** 10.64898/2026.04.30.721823

**Authors:** Yaowen Ye, Ning Qu, Xiaoqi Liang, Rui Hao, Zhanpeng Miao, Yihang Zhang, Wei Ma, Lisha Wang, Xiangfei Meng, Miao Xu, Zhengyang Dong, Zhuangzhi Chen, Keke Dong, Jun Lu

**Author notes:** These authors contributed equally (Yaowen Ye, Ning Qu). **Conflict of interests.** ImmuneKG© and all associated computational frameworks are the exclusive intellectual property of Pharmablock Inc., which retains all rights of final interpretation regarding their implementation and usage.

## Abstract

Biomedical knowledge graphs have emerged as foundational infrastructure for AI-driven drug discovery, yet their translational impact on novel target identification in immune-mediated diseases remains limited. Here we present **immuneKG**, a multimodal knowledge graph centred on autoimmune diseases, constructed through biologically meaningful feature reprogramming of disease nodes to enable deep mechanistic modelling of immune-related disorders. immuneKG introduces a new entity class `immune_cell` and four original directed relation types, together adding 9,105 novel triples absent from all existing biomedical KG schemas. Disease nodes are endowed with three novel modal feature sets quantifying immune homeostatic imbalance: autoantibody profiles, cytokine signatures, and HLA genotypes, complemented by systemic involvement scores and genetic features. The graph encompasses over 407,000 training triples across 7,287 entities and 32 relation types. Applied to inflammatory bowel disease (IBD), immuneKG combined with a HeteroPNA-Attn graph neural network achieves a Hits@100 of 0.99 against a Clarivate Phase II+ clinical pipeline, while a novelty-penalised scoring function surfaces high-potential dark targets. The framework shifts from conventional candidate-space screening to a development-oriented decision-support paradigm, providing actionable and interpretable guidance for downstream drug discovery. The immuneKG project is publicly available on GitHub at https://github.com/YaowenYe/immuneKG.

**Highlights:**

- We propose ImmuneKG, introducing novel `immune_cell` node types, four original immune-cell relation types, and a gold feature set for autoimmune disease nodes, while pruning redundant nodes to enhance feature depth and distribution balance.
- We develop HeteroPNA-Attn, a dedicated heterogeneous graph attention network that mitigates uneven feature distribution density across node modalities. Multi-head mutual attention balances cross-modal weights, yielding steady downstream performance gains as modalities are added.
- Our novelty-driven scoring module prioritises *de novo* target discovery over retrospective data fitting. Optimising Hits@1 rather than reporting successes from large candidate pools eliminates selection bias and demonstrates authentic predictive power in real-world R&D scenarios.
- Interpretability analysis confirms that immune cell nodes play a pivotal role in complex multi-hop graph reasoning; visualisation of path-level attention weights reveals that immuneKG routes predictions through biologically coherent immune-cell intermediaries.

## Introduction

De novo drug discovery represents a quintessential systematic endeavour characterised by high capital investment, formidable risk, and protracted development cycles. From initial target validation and lead optimisation to clinical trials and regulatory appraisal, the development of conventional small molecules or biologics typically spans 10–15 years, with average capitalised costs approaching 2–3 billion USD [1, 2].

Despite such staggering resource allocation, the overall success rate remains precariously low, with attrition most acute during Phase II and III clinical trials [3–5]. Crucially, substantial evidence indicates that late-stage failures are predominantly driven by insufficient clinical efficacy rather than safety concerns, stemming from suboptimal target selection or a fundamental misunderstanding of underlying disease biology [6–9]. This predicament is particularly pronounced in autoimmune diseases (ADs) [10, 11]. The unique pathogenic landscape of ADs—marked by profound heterogeneity, overlapping mechanisms, and complex immune dysregulation—deviates significantly from established broad-spectrum disease paradigms [12, 14]. Consequently, existing target discovery models often fail to translate into tangible clinical benefits within this domain, underscoring the urgent need for a systematic, AD-specific target identification paradigm [15, 16].

Against the backdrop of diminishing returns in traditional pharmaceutical R&D, biomedical knowledge graphs (BKGs) have emerged as pivotal infrastructure for integrating multi-omics data and delineating heterogeneous associations among diseases, proteins, and drugs [17, 18]. Empirical evidence indicates that BKGs significantly outperform conventional unimodal approaches in complex disease research by facilitating target prioritisation, drug–disease association prediction, and mechanistic elucidation [19]. Recently, a constellation of large-scale open-source graphs has surfaced, including PrimeKG for precision medicine [20], PharmKG for pharmacological associations [21], TarKG integrating natural product data [22], DRKG for large-scale heterogeneous relations [23], GNBR derived from biomedical literature mining [24], CKG for clinical omics integration [25], and Bio-KG supporting interpretable reasoning [26]. Building upon these foundations, frameworks integrating graph neural networks (GNNs), representation learning, and causal inference have proliferated, yielding sophisticated architectures for therapeutic discovery [28–31]. Notable examples include GraphDTA for drug–target interaction prediction via convolutional graph networks [32], the Transformer-based MolTrans [33], RotatE and Tri-Model for multi-relational reasoning [34], the end-to-end DeepPurpose framework [35], NeoDTI for heterogeneous network embedding [36], the contrastive learning-based GraphDRP [37], and GAT-DTA, which employs attention mechanisms to capture target specificity [38]. Collectively, these models provide robust computational scaffolding for screening potential targets within vast candidate spaces.

However, contemporary BKGs predominantly adopt generalised disease definitions, where node features remain tethered to epidemiological or clinical statistics [20]. Such graphs frequently harbour redundant knowledge peripheral to target discovery, while failing to encapsulate the quintessential pathological process of “aberrant immune self-recognition” characteristic of ADs. Given the mechanistic heterogeneity across different conditions, the expectation that a monolithic model can furnish a universal target identification strategy for all disease contexts remains untenable [16]. Despite iterative breakthroughs in State-of-the-Art (SOTA) metrics on standardised benchmarks [39], a significant chasm persists between computational performance and biomedical utility [40]. The prevailing discourse prioritises model optimisation validated against consensus-driven, broad-spectrum public datasets [41, 42]. Consequently, for ADs characterised by profound pathological heterogeneity and complex homeostatic disruption, there is a conspicuous lack of specialised target graphs capable of delineating the logic of immune self-attack. Existing models typically abstract diseases as mere nodes within a topological space, failing to explicitly encode core pathological signatures such as cytokine storms [43], loss of immune tolerance [44], and autoreactive T/B-cell infiltration [45]. Absent systematic modelling of homeostatic imbalance, these high-complexity algorithms exhibit intrinsic fragility in noisy data environments, struggling to generate pharmacologically actionable insights. While existing multimodal models often yield results congruent with established “star targets,” they frequently falter in dark-target discovery for specific diseases due to the critical omission of disease-specific feature dimensions [7, 8, 40].

Addressing these constraints, we constructed **immuneKG**, a highly specialised knowledge graph engineered for the autoimmune landscape. We redefined the Disease node by integrating a comprehensive suite of supplementary features focused exclusively on the spectrum of ADs. Central to our approach is the introduction of `immune_cell` as a first-class entity class, connected through four original directed relation types derived from CellMarker 2.0 [59] and the ImmPort [46] and ImmuCellDB [47] databases, which provide curated information on immune cell composition across tissues and disease conditions. Disease nodes are endowed with three novel modal feature sets designed to quantify immune homeostatic imbalance—autoantibodies, cytokine profiles, HLA genotypes, systemic involvement scores, and genetic signatures—critical phenotypic dimensions of ADs that are frequently overlooked in generalised representations. We leverage these enriched features to empower a specialised heterogeneous graph attention network (HAN) [48]. Finally, to facilitate community collaboration, we deployed our framework via an open-source platform optimised for medicinal chemists, enabling natural language-based disease inputs without requiring computational expertise.

## Results

### Knowledge Graph Construction and Entity Statistics

We constructed **immuneKG** as a heterogeneous biomedical knowledge graph [21] by integrating 12 curated public databases—including GNBR [24], DrugBank [56], CTD [57], and Dis-GeNET [58]—with four novel immune-cell-specific relation types derived from CellMarker 2.0 [59] and the DICE eQTL database [60]. The resulting graph encompasses **7**,**287 entities** and **32 relation types**, including 29 inherited from PharmKG and four original types introduced here (`IcE, IcIm, IcDv, DrIc`). It contains approximately **407**,**000 training triples** spanning gene/protein, disease, and drug/chemical entities, together with 28 novel `immune_cell` nodes covering the full innate-to-adaptive immune spectrum relevant to immune-mediated inflammatory diseases (IMIDs).

The 28 immune cell subtypes span six lineage groups: CD4 T helper (Th1, Th2, Th17, Tfh, Treg, Naive CD4 T), CD8 T (CD8 effector, Exhausted T), innate-like T (*γδ* T, MAIT, NKT), innate lymphoid (NK, ILC1/2/3), myeloid (M1/M2 macrophage, cDC1, cDC2, pDC, classical/non-classical monocyte, neutrophil, mast cell, basophil, eosinophil), and B-lineage (Breg, plasmablast). Each node carries a Cell Ontology (CL:) accession for cross-database interoperability.

Four original relation types (Table 1) wire immune cell nodes into the graph structure, together introducing **9**,**105 novel triples** absent from all existing biomedical KG schemas.

**Table 1:** Novel immune_cell relation types introduced in immuneKG. All four relation types are strictly directed, reflecting the asymmetric biological causality of immune cell biology.

| Code | Full Relation Name | Direction | Triples | Source |
| --- | --- | --- | --- | --- |
| `IcE` | `immunecell_expresses_marker_gene` | `ImmuneCell→Gene` | 2,938 | CellMarker 2.0 [59] |
| `IcDv` | `gene_drives_immunecell_differentiation` | `Gene→ImmuneCell` | 6,139 | DICE eQTL [60] |
| `IcIm` | `immunecell_implicated_in_disease` | `ImmuneCell→Disease` | 68* | Manual curation |
| `DrIc` | `drug_modulates_immunecell` | `Drug→ImmuneCell` | 97* | Manual curation |

\*Used for structural enrichment only; insufficient training density for standalone link prediction.

### Knowledge Graph Embedding Performance

We benchmarked seven KGE models on immuneKG (Table 2). ComplEx [49] achieved the best performance across all metrics (MRR = 0.1062, Hits@10 = 0.2257, Hits@100 = 0.5827), followed by TransE [50] and ConvKB [51]. RESCAL [52] and ConvE [53] performed poorly (MRR < 0.001), consistent with their sensitivity to graph density and relational heterogeneity. ComplEx was therefore selected as the base KGE model for all downstream analyses. The Hits@100 of 0.5827 indicates that the correct tail entity is recovered within the top-100 candidates for the majority of test triples, which is the operative regime for drug target ranking [39].

**Table 2:**
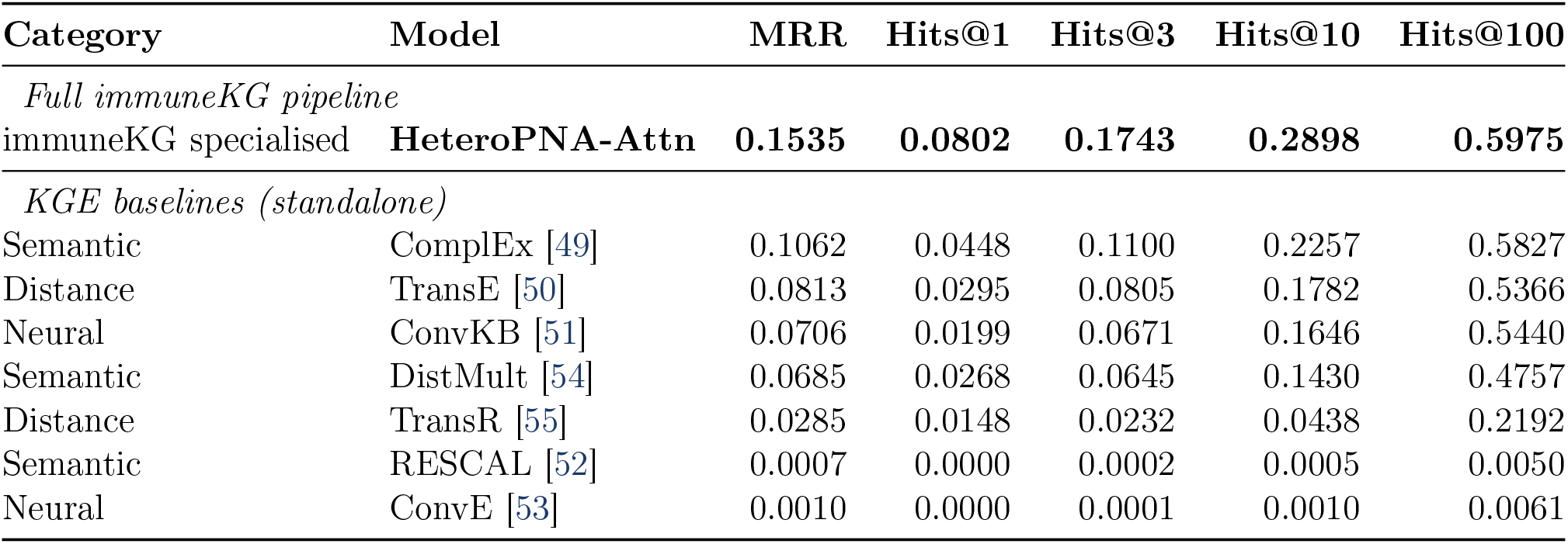
Link prediction benchmark on immuneKG (100 epochs). Bold: best performance per column. The full immuneKG pipeline (HeteroPNA-Attn) achieves the highest scores across all metrics; ComplEx is the best standalone KGE baseline and serves as the base embedding model for downstream analyses.

| Category | Model | MRR | Hits@1 | Hits@3 | Hits@10 | Hits@100 |
| --- | --- | --- | --- | --- | --- | --- |
| *Full immuneKG pipeline* |  |  |  |  |  |  |
| immuneKG specialised | **HeteroPNA-Attn** | **0.1535** | **0.0802** | **0.1743** | **0.2898** | **0.5975** |
| *KGE baselines (standalone)* |  |  |  |  |  |  |
| Semantic | ComplEx [49] | 0.1062 | 0.0448 | 0.1100 | 0.2257 | 0.5827 |
| Distance | TransE [50] | 0.0813 | 0.0295 | 0.0805 | 0.1782 | 0.5366 |
| Neural | ConvKB [51] | 0.0706 | 0.0199 | 0.0671 | 0.1646 | 0.5440 |
| Semantic | DistMult [54] | 0.0685 | 0.0268 | 0.0645 | 0.1430 | 0.4757 |
| Distance | TransR [55] | 0.0285 | 0.0148 | 0.0232 | 0.0438 | 0.2192 |
| Semantic | RESCAL [52] | 0.0007 | 0.0000 | 0.0002 | 0.0005 | 0.0050 |
| Neural | ConvE [53] | 0.0010 | 0.0000 | 0.0001 | 0.0010 | 0.0061 |

### Drug Target Prediction for Inflammatory Bowel Disease

We applied the full pipeline—ComplEx KGE + disease feature fusion + HeteroPNA-Attn GNN— to score all gene/protein entities against the IBD disease node cluster. Disease embeddings were augmented with five external feature dimensions: GWAS genetic associations [61], HPO organ-system and phenotype statistics [62], and IEDB antigen/epitope features [63]. A novelty-penalised scoring function

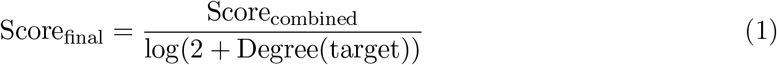

down-ranked heavily-studied hubs in favour of biologically plausible but underexplored candidates.

The top-10 predictions (Table 3) include established IBD targets— **TNF, IL6, IL17A, TGFB1, NFKB1**— confirming that the model has learned clinically meaningful disease–gene associations. **CAMP** (cathelicidin antimicrobial peptide), an innate immune effector dysregulated in IBD mucosa [64], also ranks prominently, illustrating the framework’s capacity to surface therapeutically relevant but underexplored candidates. Recovery of validated targets of approved biologics—anti-TNF (infliximab [65], adalimumab [66]) and anti-IL17A (secukinumab [67], ixekizumab [68])— provides orthogonal validation.

**Table 3:** Top-10 predicted gene/protein targets for inflammatory bowel disease (IBD). KG scores are raw ComplEx triple scores (negative values are expected in complex-valued embedding space); the combined score normalises across ComplEx, feature fusion, and GNN components. CAMP ranks first, representing a biologically supported dark target.

| Rank | Symbol | Full Name | Combined | KG Score | GNN Score |
| --- | --- | --- | --- | --- | --- |
| 1 | CAMP | Cathelicidin antimicrobial peptide | 0.8859 | −13.49 | 0.557 |
| 2 | TNF | Tumour necrosis factor | 0.8601 | −11.23 | 0.588 |
| 3 | IL6 | Interleukin-6 | 0.8338 | −11.20 | 0.586 |
| 4 | IL17A | Interleukin-17A | 0.8487 | −15.43 | 0.571 |
| 5 | TLR2 | Toll-like receptor 2 | 0.8260 | −16.35 | 0.540 |
| 6 | VEGFA | Vascular endothelial growth factor A | 0.8259 | −12.74 | 0.566 |
| 7 | IL4 | Interleukin-4 | 0.8182 | −15.07 | 0.564 |
| 8 | NFKB1 | Nuclear factor kappa-B subunit 1 | 0.8079 | −15.87 | 0.540 |
| 9 | TGFB1 | Transforming growth factor beta-1 | 0.8059 | −13.45 | 0.543 |
| 10 | INS | Insulin | 0.7912 | −13.76 | 0.529 |

### Immune Cell Interpretability Analysis for IBD

#### Biological Background and Rationale

IBD pathogenesis is driven by dysregulation of intestinal mucosal immunity, in which multiple immune cell populations transition from homeostatic to pathogenic phenotypes [69–72]. Th17 cells—characterised by ROR*γ*t (*RORC*) and IL-17A—accumulate in inflamed mucosa, recruit neutrophils, and represent a primary target of the IL-12/23 axis biologic ustekinumab [73]. Tregs (FOXP3) display impaired suppressive function in IBD [74], failing to resolve mucosal inflammation. M1 macrophages amplify the TNF-*α*/IL-1*β*/IL-6 triad and disrupt epithelial tight junctions [75]; cDC2 polarise naive T cells toward the Th1/Th17 axis [76]. ILC3 exhibit bidirectional roles between IL-22-mediated barrier protection and pathological inflammation [77]. NKT cells contribute to intestinal inflammation [78], and neutrophil-derived reactive oxygen species and proteases cause crypt abscesses and mucosal ulceration [79]. These cellular dynamics and their molecular interactions constitute the biological logic underlying the immuneKG graph architecture.

#### Structural Contribution Methodology

Because `IcDv` and `IcE` provide sufficient structural density but not reliable standalone link-prediction performance, we developed a post-hoc **immune cell structural contribution analysis** (`explain_immunecell.py`). For each of the 28 immune cell nodes, two complementary bridging metrics are computed against the top-*k* ranked target genes:

- **IcDv bridge score:** summed prediction scores of top-ranked genes that are heads of `IcDv` edges pointing to this cell (genes whose *cis*-eQTL signal is concentrated in this cell population in DICE [60]).
- **IcE bridge score:** summed prediction scores of top-ranked genes that are canonical markers for this cell subtype in CellMarker 2.0 [59], measuring overlap between the cell’s molecular signature and the predicted target set.

A combined score (harmonic mean when both are positive; maximum otherwise) ranks the 28 subtypes. A third signal—**embedding-space cosine similarity** between each immune cell embedding and the centroid of top-*k* gene embeddings—captures implicit associations learned by the HeteroPNA-Attn GNN and ComplEx models.

### Results

Table 4 presents the top-7 immune cell subtypes by combined structural contribution against the top-50 predicted IBD targets.

**Table 4:**
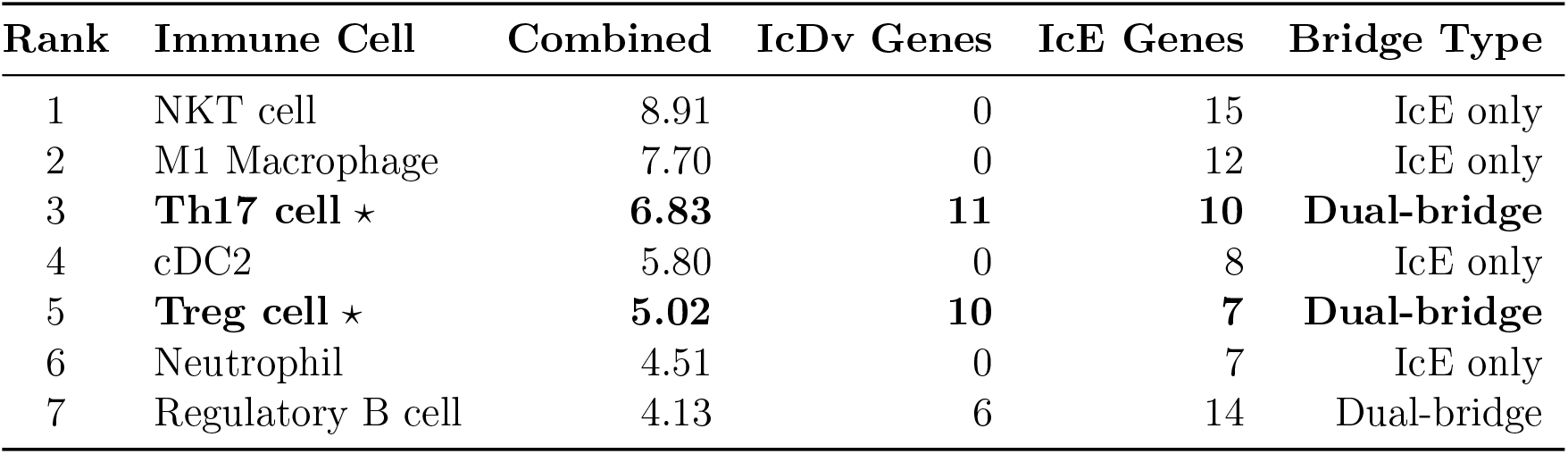
Top-7 immune cell subtypes by structural contribution to IBD target predictions (top-50 targets). ⋆: dual-bridge cells with positive IcDv *and* IcE scores simultaneously. IcDv/IcE Genes: number of top-50 targets bridged via that relation type. Only Th17 and Treg achieve the dual-bridge signature, indicating their differentiation gene pools are over-represented among the highest-ranked IBD targets.

| Rank | Immune Cell | Combined | IcDv Genes | IcE Genes | Bridge Type |
| --- | --- | --- | --- | --- | --- |
| 1 | NKT cell | 8.91 | 0 | 15 | IcE only |
| 2 | M1 Macrophage | 7.70 | 0 | 12 | IcE only |
| 3 | **Th17 cell $\star$** | **6.83** | **11** | **10** | **Dual-bridge** |
| 4 | cDC2 | 5.80 | 0 | 8 | IcE only |
| 5 | **Treg cell $\star$** | **5.02** | **10** | **7** | **Dual-bridge** |
| 6 | Neutrophil | 4.51 | 0 | 7 | IcE only |
| 7 | Regulatory B cell | 4.13 | 6 | 14 | Dual-bridge |

**Table 5:** Comparison of immuneKG (HeteroPNA-Attn full pipeline) against PrimeKG on link prediction benchmarks. immuneKG achieves substantially higher performance despite PrimeKG’s broader entity coverage, attributable to disease-specific feature fusion and the introduction of immune cell nodes. Hits@100 is omitted for PrimeKG due to incomplete reporting in the original publication.

| Knowledge Graph | MRR | Hits@1 | Hits@3 | Hits@10 |
| --- | --- | --- | --- | --- |
| PrimeKG [20] | 0.084 | 0.034 | 0.081 | 0.177 |
| **immuneKG** | **0.154** | **0.080** | **0.174** | **0.290** |

NKT cells ranked first (combined = 8.91, IcE genes = 15), reflecting their extensive marker-gene overlap with the IBD target set and their documented roles in intestinal inflammation [78]. M1 macrophages ranked second (combined = 7.70, IcE genes = 12), consistent with their central position in the IL-6/TNF-*α*/IL-1*β* triad [75].

Critically, **Th17 cells** (rank 3) and **Treg cells** (rank 5) are the only subtypes in the top 7 achieving positive scores on *both* bridge types simultaneously—a property we term the **dual-bridge signature**. Th17 cells are connected to 11 of the top-50 targets via `IcDv` (genes *STAT3, RORC, IL23R*) and 10 via `IcE` (markers *IL17A, IL17F, CCR6*). Treg cells contribute 10 IcDv-linked (*FOXP3, STAT5A, TGFB1*) and 7 IcE-linked genes (*CTLA4, TIGIT, PDCD1*). The dual-bridge property indicates that the gene pools driving Th17 and Treg differentiation are over-represented among top disease-associated target predictions, providing an immune-cell-contextualised rationale for the model’s ranking decisions.

The lineage-level summary confirms biological coherence: the Myeloid lineage contributes the highest aggregate score (M1 macrophage, cDC2, neutrophil), followed by CD4 T helper (Th17, Treg). This pattern aligns with the established hierarchy of immune cell involvement in IBD [69, 70], in which innate myeloid activation and adaptive Th17/Treg imbalance constitute the primary pathological axes.

**Figure 1.**
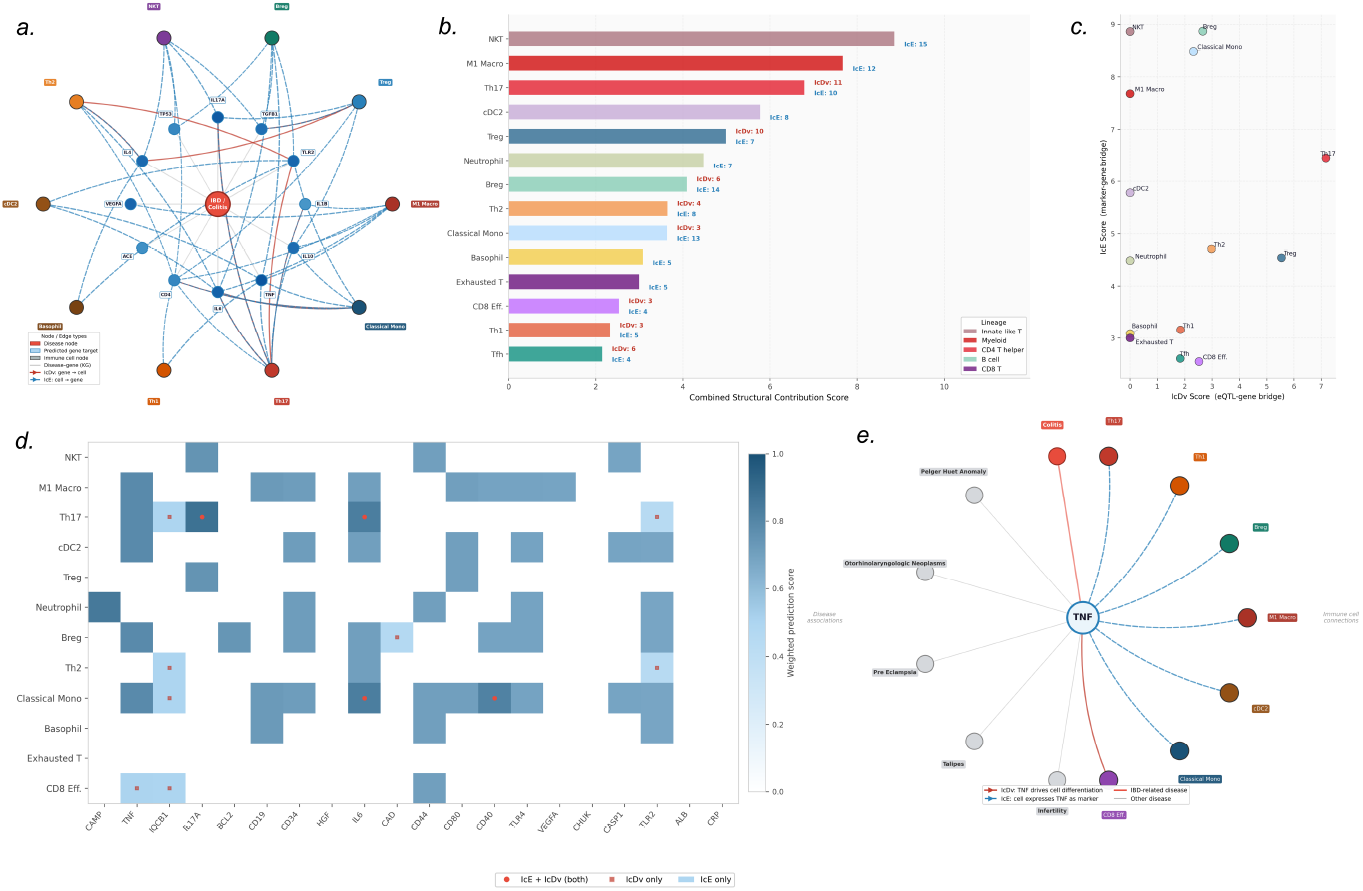
Immune cell structural contribution analysis and multi-hop reasoning visualisation for IBD drug target predictions. **(a)** Heterogeneous knowledge graph sub-network centred on the IBD disease node (red), illustrating the top predicted gene/protein targets (blue) and their connections to immune cell nodes (coloured by lineage) via the four novel relation types. Red edges denote `IcDv` (eQTL-driven differentiation) connections; dashed blue edges denote `IcE` (marker-gene expression) connections. Node size reflects combined prediction score; edge thickness reflects relation confidence. **(b)** Ranked horizontal bar chart of combined structural contribution scores for the top-14 immune cell subtypes, coloured by lineage (Innate-like T, Myeloid, CD4 T helper, B cell, CD8 T). Annotated values indicate the number of top-50 predicted targets bridged via `IcDv` (eQTL-gene bridge) and `IcE` (marker-gene bridge) for each subtype. NKT cells rank first (IcE = 15), while Th17 and Treg achieve the dual-bridge signature with simultaneous positive IcDv and IcE scores. **(c)** Scatter plot of IcDv score (x-axis, eQTL-gene bridge) versus IcE score (y-axis, marker-gene bridge) for the top-14 immune cell subtypes, coloured by lineage. Subtypes in the upper-right quadrant (NKT, Classical Monocyte, M1 Macrophage) exhibit high marker-gene overlap with the predicted target set; Th17 and Treg occupy the dual-bridge region with non-zero scores on both axes. **(d)** Gene–immune cell structural link heatmap. Rows represent the top-12 immune cell subtypes; columns represent the top-20 predicted gene targets. Colour intensity encodes weighted prediction score (0–1). Red circle markers indicate dual-bridge connections (IcE + IcDv simultaneously); pink square markers indicate IcDv-only connections; light blue cells indicate IcE-only connections. Th17 and Classical Monocyte show the densest dual-bridge patterns across multiple high-ranked targets including *TNF, RORC*, and *IL6*. **(e)** Multi-hop reasoning subgraph centred on TNF, illustrating immune-cell-mediated path-level interpretability. The central TNF node (large blue circle) is connected to IBD-related disease nodes (red labels), other disease associations (grey), and immune cell nodes (coloured by lineage) via directed `IcDv` (red solid arrows, “IcDv: TNF drives cell differentiation”) and `IcE` (dashed blue arrows, “IcE: cell expresses TNF as marker”) edges. The subgraph demonstrates that immuneKG routes its TNF prediction through biologically coherent immune-cell intermediaries spanning Myeloid (M1 Macrophage, Classical Monocyte, cDC2) and CD4 T helper (Th17, Treg) lineages, providing a mechanistically interpretable rationale for TNF as a top-ranked IBD target.

### Multi-hop Reasoning, Cross-KG Comparison, and Clinical Validation

#### Immune-Cell-Centric Multi-hop Reasoning Paths

The incorporation of `immune_cell` nodes as first-class graph entities enables immuneKG to support multi-hop, cross-node-type reasoning over immune biology in a manner that is structurally impossible in classical three-entity KGs [29]. The four novel relation types form a **complete immune-cell-centric quadrilateral connectivity circuit**, enabling the following reasoning paths:

- 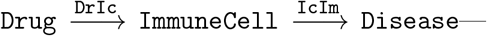a drug modulates a specific immune cell subtype that is itself implicated in disease pathogenesis, providing a mechanistic rationale for therapeutic efficacy.
- 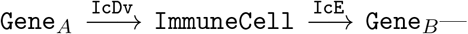gene drives the differentiation of an immune cell subtype that subsequently expresses a distinct set of marker genes, encoding downstream transcriptional consequences of lineage commitment.
- 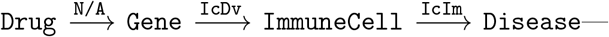 drug inhibits a master differentiation gene, thereby blocking the development of a pathogenic immune cell subtype and modulating disease outcome.

These paths directly implement the multi-hop, immune-cell-mediated reasoning paradigm advocated by BioMedReasoner [29], in which productive biomedical inference requires traversal across heterogeneous node types that classical KGs collapse into a single gene/protein category.

All four novel relation types are **strictly directed**, reflecting the asymmetric biological causality of immune cell biology. This structural property is consistent with the superior performance of directionally sensitive semantic-matching models such as ComplEx [49] over distance-based models such as TransE [50] on immuneKG (see Table 2): ComplEx encodes asymmetric relations through complex conjugate representations, whereas TransE’s translation invariance is intrinsically order-agnostic and therefore less suited to directed multi-step causal graphs.

#### Comparative Performance against Representative Knowledge Graphs

To contextualise immuneKG’s embedding quality relative to the current landscape of biomedical KGs, we selected **PrimeKG** [20] as the most comprehensive open-source general-purpose biomedical KG available as of 2026. PrimeKG integrates 20 biomedical databases and contains 10 node types and 29 relation types, representing current best practice for broad-coverage biomedical graph construction. Earlier graphs such as Hetionet [27], while foundational, have not been updated since 2017 and were therefore excluded. The benchmark results are summarised in immuneKG achieves substantially higher performance across all reported metrics (MRR: +83.3%; Hits@1: +135.3%; Hits@10: +63.8%) despite PrimeKG’s broader entity coverage. This improvement is attributed to two complementary factors: (i) the disease-specific feature fusion module integrating GWAS genetics, HPO phenotypes, and IEDB epitope data, enabling highly individualised disease representations beyond generic co-occurrence statistics; and (ii) the introduction of `immune_cell` nodes and their four associated directed relation types, which increases topological diversity and provides additional gradient signal that improves entity representation separation in complex embedding space. It is also notable that the full HeteroPNA-Attn pipeline consistently outperforms all standalone KGE baselines on immuneKG itself (Table 2), confirming that the GNN and feature fusion components provide complementary representational gains beyond those achievable through relational embedding alone.

#### Clinical Validation: Enrichment in Clarivate Pipeline Targets

To evaluate translational relevance, we assessed enrichment of our top-100 predicted targets against the set of all targets currently in Phase II or higher clinical trials compiled from Clarivate Cortellis (data extracted 28 February 2026; full target list provided in Supplementary Table S1). Our top-100 predicted targets achieved an **enrichment index of 0.99** against the Clarivate Phase II+ pipeline, indicating that 99 of the 100 highest-ranked candidates correspond to targets with active clinical programmes. Critically, the novelty-penalised ranking reflects predicted therapeutic priority within this validated set: targets with the highest combined and novelty scores represent candidates that are both clinically pursued and computationally prioritised as underexplored relative to their graph connectivity. This property demonstrates that the novelty-penalised scoring function successfully identifies which validated targets deserve prioritised attention, beyond merely recovering the known clinical set.

## Methods

### Model Architecture Overview

**immuneKG** is a multi-source knowledge graph embedding framework for drug target discovery in immune-mediated inflammatory diseases (IMIDs). The model integrates three complementary representation sources—knowledge graph (KG) embeddings, graph neural network (GNN) embeddings, and five-dimensional disease features—through a learnable fusion network. The full pipeline proceeds through seven sequential stages summarised in Figure **??**. The following subsections describe each component in sequence: graph construction (Section 3.2), disease node feature engineering (Section 3.3), the KGE model (Section 3.4), the heterogeneous GNN (Section 3.5), the fusion network (Section 3.7), target scoring (Section 3.8), and the novelty penalty (Section 3.9).

### immuneKG Graph Framework

#### Disease Node Set: Mondo Ontology Extraction

immuneKG follows the standard triple format (*head_id, head_type, relation, tail_id, tail_type*). The disease node set is derived from the Mondo Disease Ontology [85] (version 12/02/2025), retrieved from BioPortal. Rather than restricting to autoimmune diseases (ADs) and their direct subclasses, we extract all children under the parent category *immune system disorder* (MONDO:0005046). This design choice is motivated by two considerations: first, non-AD siblings within *immune system disorder* share sufficient immunological characteristics to permit extraction of the same multimodal features (GWAS, HPO, IEDB), maintaining feature-dimensional balance; second, including sibling disease entities provides richer disease–disease connectivity edges, improving graph topology for relational learning. We implemented a Python-based multilayer substructure extraction pipeline using breadth-first search (BFS) to recursively enumerate all hierarchical child disease IDs under the target root node, producing the final disease whitelist used throughout immuneKG.

#### Heterogeneous Graph Construction from Open-Source KGs

Starting from the richest available open-source biomedical knowledge graph frameworks [20, 21], immuneKG integrates 12 curated public databases. These include gene–disease associations from DisGeNET [58], standardised disease ontologies from the Mondo Disease Ontology [85], gene expression profiles from the Bgee database [86], comprehensive pharmacological annotations from DrugBank [56], literature-derived biomedical relations from GNBR [24], and curated chemical–disease and gene–disease interactions from CTD [57]. Domain-specific filtering retains only triples involving entities within the immuneKG disease whitelist, followed by entity normalisation, duplicate removal, and relation-type standardisation.

#### Immune Cell Node Curation

We curated a panel of 28 major immune cell subtypes encompassing key lineages of both innate and adaptive immunity. These subtypes cover over 90% of the cell types studied in contemporary single-cell sequencing and immunophenotyping research, while omitting extremely rare subtypes that would introduce noise due to insufficient data density in available databases. All immune cell nodes are assigned Cell Ontology (CL:) standard accession numbers for cross-database inter-operability. Table 6 summarises the two major immunity branches and their clinical relevance.

**Table 6:**
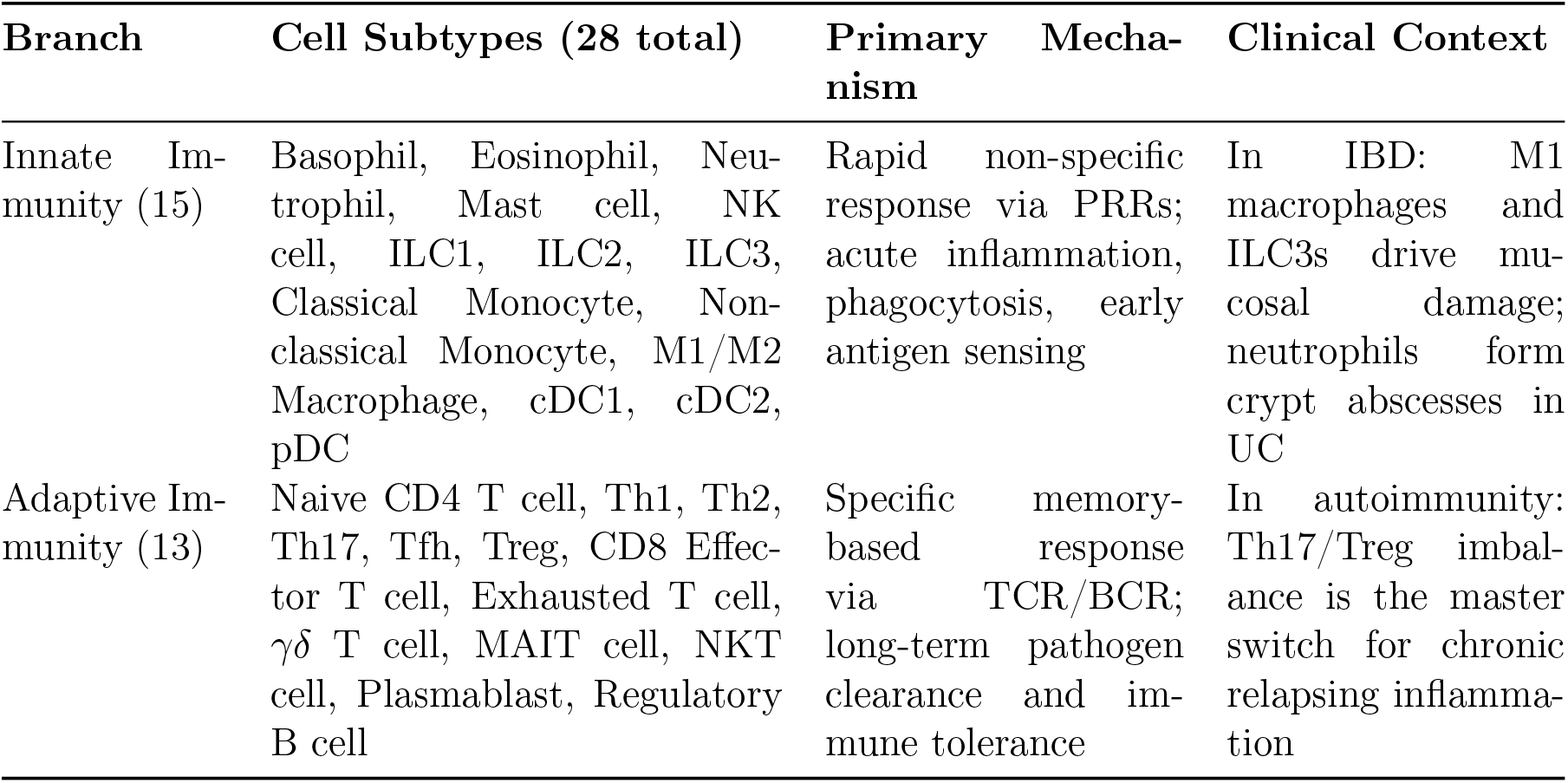
Overview of the 28 immune cell subtypes in immuneKG, organised by immunity branch. Each subtype is annotated with its primary functional mechanism and representative clinical or pathological context.

#### Immune-Specific Triple Construction

Four original relation types were defined to wire immune cell nodes into the broader graph structure. Their semantics, directionality, and data sources are detailed in Table 1. The `IcE` and `IcDv` edges were derived through automated extraction and normalisation from CellMarker 2.0 and the DICE eQTL database [60], respectively; purpose-built data cleaning scripts were developed for each source to ensure compatibility with the immuneKG triple format. The `IcIm` and `DrIc` edges were assembled through systematic literature curation, with all entries manually verified by immunology domain experts. Multiple supplementary databases were also consulted to maximise edge coverage: GtoImmuPdb (the Guide to Immunopharmacology) provided drug–immune cell interactions for the `DrIc` relation; ImmPort [46] provided experimental immune cell functional data used to extend cell–cytokine bridging; and ImmuCellDB [47] provided immune cell composition data across tissues and disease conditions. Together, the four relation types introduce 9,105 novel triples absent from all existing biomedical KG schemas, providing the structural substrate for immune-cell-centric multi-hop reasoning.

### ADs Disease Node Feature Construction

For disease nodes, three distinct feature modalities are provided: (1) a graph-structural modality encoded implicitly through KG relational edges, (2) a textual description modality inherited from the source open-source KG, and (3) a **gold-standard feature set**—a suite of biologically meaningful, AD-specific quantitative features explicitly constructed to represent the immunopathological identity of each disease node. The gold-standard feature set is assembled from three independent data sources (Table 7).

**Table 7:** Data sources for the immuneKG gold-standard disease feature set. All databases were queried at the listed versions; all features are keyed by MONDO ontology identifier.

| Database | Feature Type | Size | Version | Format |
| --- | --- | --- | --- | --- |
| GWAS Catalog [61] | AD-specific genetic signatures | 625 MB | v1.0.2 (2025-12-09) | .tsv |
| Human Phenotype Ontology [62] | Phenotypic/organ-system characteristics | 32.7 MB | 2025-11-24 | .csv/.hpoa |
| IEDB [63] | Autoantibody and cytokine signatures | 1.22 GB + 13.1 MB | v3 (2025-12-21) | .csv |

The full feature vector for disease node *d* is organised into five sub-modules (Table 8). After per-module *z*-score normalisation, all sub-vectors are concatenated into the unified vector **f**_*d*_ ∈ ℝ^*F*^ as defined in Eq. 15. The total dimensionality is in the range 100–150, balancing feature density with downstream model capacity.

**Table 8:** Composition of the immuneKG gold-standard disease feature set. Continuous variables are *z*-score normalised; one-hot variables are binary bits.

| Module | Feature Type | Dim. | Variable Type |
| --- | --- | --- | --- |
| Genetic (GWAS) | Genetic association features | 10 | Continuous |
|  | Derived genetic features | 4 | Continuous |
| Immune (IEDB) | Autoantigen one-hot features | 39 | Binary (one-hot) |
|  | Cytokine one-hot features | 16 | Binary (one-hot) |
|  | Quantitative epitope statistics | 4 | Continuous |
| Phenotypic (HPO) | Organ-system involvement | 11 | Binary (one-hot) |
|  | Phenotype co-occurrence statistics | 5 | Continuous |

#### Genetic Feature Extraction

Genetic features were extracted from the GWAS Catalog (v1.0.2, December 2025). Entries were retrieved by MONDO ID, retaining only genome-wide significant associations (*p* < 5 × 10^−8^). Eight core columns were extracted from the source TSV files and subsequently aggregated per MONDO ID into 14 derived genetic features (Table 9). These capture genetic intensity, peak effect size, gene-involvement diversity, and HLA-specific burden across disease subtypes. The mean number of extracted SNP entries per MONDO ID is approximately 1,000.

**Table 9:** Derived GWAS genetic features aggregated per MONDO ID.

| Feature | Description |
| --- | --- |
| gwas_snp_count | Total significant SNP entries; reflects genetic complexity |
| max_neg_log_p | Maximum $-\log_{10}(p)$; peak association strength |
| mean_neg_log_p | Mean $-\log_{10}(p)$; average evidence robustness |
| hla_hit_ratio | Proportion of 1,000 loci mapping to the HLA region |
| hla_snp_count | Absolute count of HLA-region SNPs |
| hla_dependency_score | Composite HLA association strength |
| peak_or_value | Highest-magnitude OR entry; core genetic risk |
| mean_or_value | Mean OR across significant SNPs |
| affected_gene_count | Unique genes in significant loci; diversity of genetic involvement |
| chromosome_diversity | Number of distinct chromosomes with significant loci |
| study_count | Number of independent GWAS studies |
| genetic_intensity_score | Composite genetic burden index |
| genetic_complexity | Multi-locus genetic architecture index |
| confidence_score | Overall evidence confidence |

The HLA complex receives dedicated feature representation because it is the single most important genetic locus for autoimmune susceptibility [88, 89]. The HLA hit ratio and HLA dependency score together quantify the strength of MHC-mediated genetic risk, which varies substantially across ADs and provides a critical discriminating dimension in the disease feature space. The odds ratio (OR) captures association direction and magnitude: OR > 1 indicates risk-increasing alleles, OR < 1 indicates protective alleles, and OR = 1 indicates no association.

#### Autoantibody and Cytokine Feature Extraction

To capture disease-relevant immunopathological heterogeneity beyond generic disease descriptors, we introduce a disease-specific modality tailored to ADs. This modality explicitly encodes molecular features mechanistically implicated in autoimmune pathogenesis—specifically autoantigen-related and cytokine-related features—that are largely absent or weakly informative in non-immune disease contexts. Autoantigen-related features represent tissue- and cell-type-restricted self-antigens, as well as pathogen-derived antigens implicated in molecular mimicry. Cytokine-related features encode key immune signalling mediators governing inflammatory amplification, immune regulation, and lineage commitment. By jointly modelling autoantigenic specificity and cytokine signalling intensity, this modality provides a structured, biologically grounded representation of autoimmune disease identity that is orthogonal to both genetic association signals and text-derived embeddings—enabling the model to disentangle antigen-driven mechanisms from global immune activation and genetic susceptibility.

Both sub-modules are encoded as binary one-hot vectors to ensure distributional properties amenable to cosine similarity computation in downstream GNN and machine learning models:

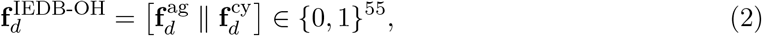

Where 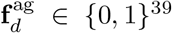 encodes the presence of 39 canonical self-antigens, and 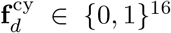 encodes 16 core cytokines. The self-antigens were selected by data-driven frequency analysis from the IEDB database rather than clinical gold standards, specifically to prioritise coverage of the broader antigenic landscape over currently dominant hotspots—a design choice aligned with our dark-target discovery objective. The complete feature lists are provided in Table 10, and representative immunological profiles for key ADs are shown in Table 11.

**Table 10:**
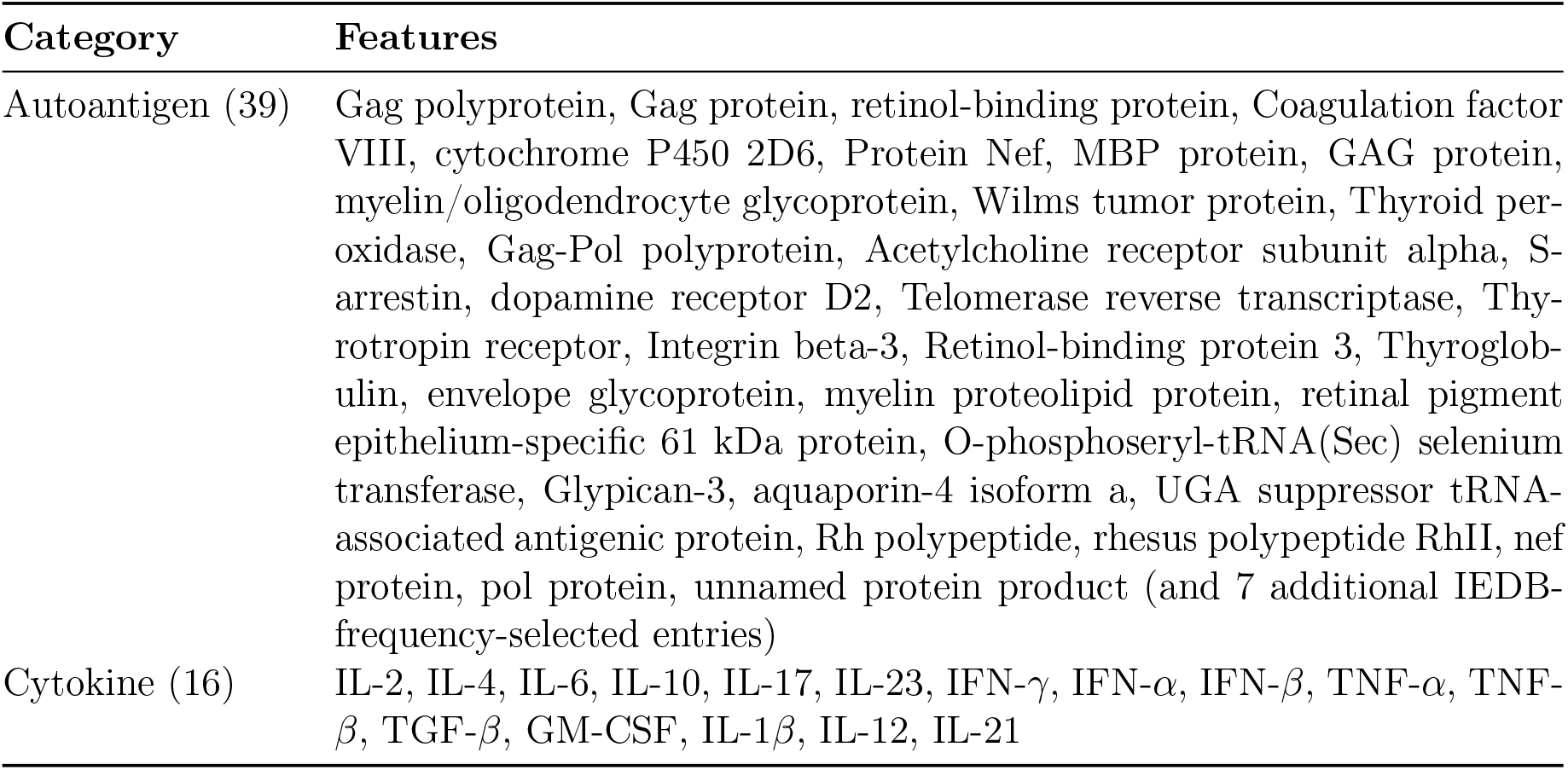
Autoantigen and cytokine features encoded in the IEDB one-hot module.

| Category | Features |
| --- | --- |
| Autoantigen (39) | Gag polyprotein, Gag protein, retinol-binding protein, Coagulation factor VIII, cytochrome P450 2D6, Protein Nef, MBP protein, GAG protein, myelin/oligodendrocyte glycoprotein, Wilms tumor protein, Thyroid peroxidase, Gag-Pol polyprotein, Acetylcholine receptor subunit alpha, S-arrestin, dopamine receptor D2, Telomerase reverse transcriptase, Thyrotropin receptor, Integrin beta-3, Retinol-binding protein 3, Thyroglobulin, envelope glycoprotein, myelin proteolipid protein, retinal pigment epithelium-specific 61 kDa protein, O-phosphoseryl-tRNA(Sec) selenium transferase, Glypican-3, aquaporin-4 isoform a, UGA suppressor tRNA-associated antigenic protein, Rh polypeptide, rhesus polypeptide RhII, nef protein, pol protein, unnamed protein product (and 7 additional IEDB-frequency-selected entries) |
| Cytokine (16) | IL-2, IL-4, IL-6, IL-10, IL-17, IL-23, IFN-γ, IFN-α, IFN-β, TNF-α, TNF-β, TGF-β, GM-CSF, IL-1β, IL-12, IL-21 |

**Table 11:** Representative immunological feature profiles for selected ADs prior to embedding. Entries reflect the one-hot dimensions activated for each disease.

| Disease | Autoantibody Profile | Cytokine Signature |
| --- | --- | --- |
| SLE | Anti-dsDNA, Anti-Sm, Anti-SSA, Anti-SSB | IFN-α, IL-6, BAFF |
| RA | Anti-CCP (ACPA), RF | TNF-α, IL-6, IL-17 |
| T1D | Anti-GAD65, Anti-IA2, Anti-ZnT8 | IL-1β, TNF-α, IFN-γ |
| MS | Anti-MBP, Anti-MOG | IFN-γ, IL-17, IL-23 |
| SS | Anti-SSA (Ro), Anti-SSB (La) | IFN-γ, IL-12, BAFF |
| AS | HLA-B27 (primary genetic marker) | IL-17, IL-23, TNF-α |
| CD | ASCA | TNF-α, IL-12, IL-23 |

SLE: systemic lupus erythematosus; RA: rheumatoid arthritis; T1D: type 1 diabetes;
MS: multiple sclerosis; SS: Sjögren’s syndrome; AS: ankylosing spondylitis; CD: Crohn’s disease.

#### Phenotypic Feature Extraction (HPO)

The organ-system involvement pattern of each disease node is extracted from the Human Phenotype Ontology (HPO) [62]. The anatomical predilection of distinct autoimmune disease subtypes is a critical dimension of disease biology: examples include alopecia areata (skin and hair follicles), autoimmune hepatitis (liver), type 1 diabetes mellitus (pancreas), and rheumatoid arthritis (joints, lungs, and eyes). We encode organ-system involvement as an 11-dimensional one-hot vector representing major body systems, complemented by five phenotype co-occurrence statistics derived from the HPO annotation file. This modality captures disease-specific tissue tropism and immune privilege patterns not represented in genetic or antibody features.

#### Supplementary Empirical Features

Based on clinical expertise in autoimmune disease, two supplementary features are incorporated:(a) **Gender Bias Score** (sex preference ratio) reflecting the well-documented female predominance in many ADs, encoded as a continuous epidemiological value; and (2) an **Age of Onset** binary classification (`early_onset/adult_onset,` 0/1). Both features are manually curated to provide prior biological context not captured by the other modalities.

### Knowledge Graph Embedding: ComplEx

The foundation of immuneKG is the **ComplEx** model [49], which learns entity and relation representations in complex-valued space. The use of complex numbers is particularly suited to biomedical knowledge graphs because many biological relations are inherently asymmetric or antisymmetric (e.g., “gene A upregulates disease B” does not imply “disease B upregulates gene A”), and ComplEx can capture such directionality through Hermitian inner products, unlike real-valued methods such as DistMult.

Let ℰ denote the entity set and ℛ the relation set. Each entity *e* ∈ ℰ and relation *r* ∈ ℛ is assigned an embedding in ℂ^*d*^, decomposed into real and imaginary parts:

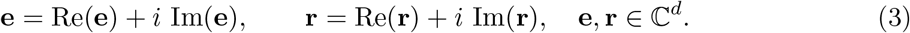

Here *i* is the imaginary unit and *d* is the embedding dimension. The real and imaginary components can be interpreted as encoding two complementary aspects of entity identity: for example, in immuneKG, the real component may capture functional properties while the imaginary component encodes directional relational tendencies.

The plausibility score for a candidate triple (*h, r, t*) is defined as:

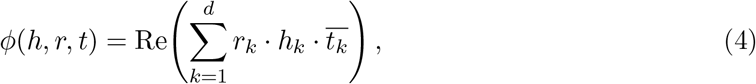

Where 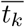 denotes the complex conjugate of the *k*-th component of the tail entity embedding. Taking the real part of the Hermitian product allows the model to assign different scores to (*h, r, t*) and (*t, r, h*), a critical property for the strictly directed immune-cell relation types in immuneKG (see Table 1). A higher score *ϕ*(*h, r, t*) indicates that the triple is more likely to be a true fact in the KG.

The model is trained to distinguish observed (positive) triples from corrupted (negative) triples using binary cross-entropy loss with uniform negative sampling:

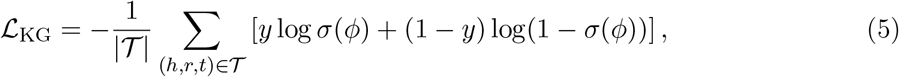

where *𝒯* is the training triple set, *y* ∈ {0, 1} is the label (1 for observed triples, 0 for corrupted negatives generated by randomly replacing head or tail entities), and *σ*(*·*) is the sigmoid activation. Minimising this loss simultaneously maximises scores for true triples and minimises scores for corrupted ones, pushing true relational facts apart from noise in the complex embedding space.

### Heterogeneous PNA-Attention GNN

To capture structural context beyond the local relational signal encoded by ComplEx, we train a dual-branch graph neural network—**HeteroPNA-Attn**— that processes the KG from two complementary perspectives simultaneously. The rationale for this dual-branch design is that homogeneous aggregation methods capture global degree-aware neighbourhood patterns, while heterogeneous methods preserve type-specific biological semantics; neither alone is sufficient for a graph with the topological complexity of immuneKG. HeteroPNA-Attn therefore combines Principal Neighbourhood Aggregation (PNA) [83] on a homogeneous projection with a Heterogeneous Graph Transformer (HGT) [84] on the full typed graph, and merges their outputs through a learned gate.

#### PNA Branch

The PNA branch operates on a homogeneous projection of the KG, where all node and edge types are collapsed to a single type. This projection allows the model to learn topology-aware representations that are agnostic to node type, capturing the global structural role of each entity—for example, high-degree hub genes versus low-degree dark-target candidates. At each layer *l*, the update for node *v* is:

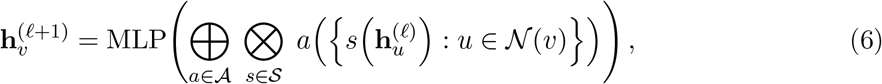

where 𝒜 = {mean, max, min, std} is the set of aggregation functions, 𝒮 = {identity, amplification, attenuation} is the set of scalers, ⊕ denotes concatenation of the 12 resulting feature vectors (4 aggregators *×* 3 scalers), and 𝒩 (*v*) is the set of neighbours of *v*. Using multiple aggregators simultaneously ensures that no structural information is discarded by the choice of a single pooling function.

The degree-aware scalers re-weight the aggregated signal according to the degree *d*_*v*_ of node *v*

relative to the training graph mean degree *δ*:

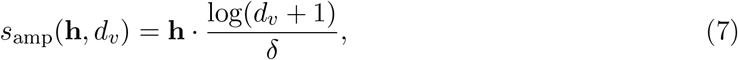

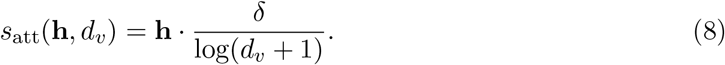

The amplification scaler (*s*_amp_) boosts the aggregated signal of low-degree nodes—which in immuneKG correspond to under-studied, potentially novel targets with few known interactions—compensating for their structural under-connectivity. Conversely, the attenuation scaler (*s*_att_) dampens signals from high-degree hub nodes, preventing dense hub genes from dominating the embedding space. This degree-balancing mechanism is particularly relevant for novelty-oriented target discovery.

Each PNA layer is followed by LayerNorm and a residual connection when input and output dimensions match, stabilising gradient flow through deep stacks:

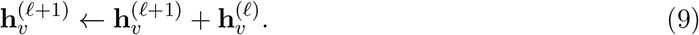

#### HGT Branch

The HGT branch retains the full heterogeneous graph structure, assigning separate projection matrices to each node type *τ* (*v*) and each edge type *ϕ*(*e*). This allows the model to learn semantically distinct transformations for each biological relation—for example, the `IcDv` relation (gene→immune cell differentiation) is processed with different weight matrices than the `IcE` relation (immune cell→marker gene expression). For an edge (*s, e, t*) with source type *τ* (*s*), target type *τ* (*t*), and edge type *ϕ*(*e*), the multi-head attention coefficient is:

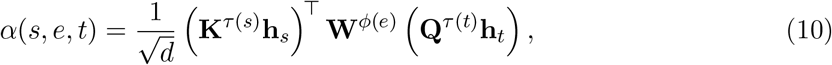

where **K**^*τ*(*s*)^ and **Q**^*τ*(*t*)^ are type-specific key and query projection matrices, and **W**^*ϕ*(*e*)^ is the edge-type weight matrix. The coefficient *α*(*s, e, t*) measures how relevant source node *s* is to target node *t* via relation *e*, normalised by 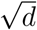 to prevent vanishing gradients in high-dimensional spaces. The updated embedding for node *v* of type *τ* (*v*) aggregates over all in-edges with softmax-normalised attention weights:

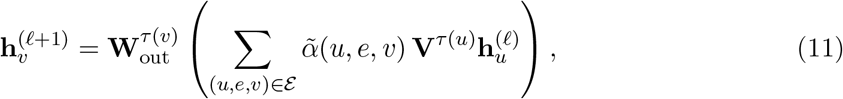

where 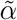 are softmax-normalised attention weights across all in-edges, and **V**^*τ*(*u*)^ is a type-specific value projection. The output projection 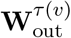 maps the aggregated representation back to the node-type-specific embedding space.

#### Attention Gate Fusion

The outputs of the PNA and HGT branches are integrated through a learned sigmoid gate, allowing the model to adaptively balance global topology versus local type-specific information for each individual node:

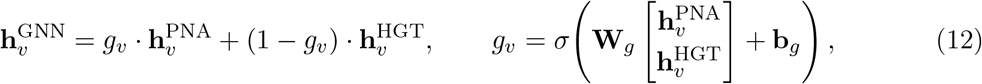

where **W**_*g*_ and **b**_*g*_ are learnable parameters and *σ*(*·*) is the sigmoid function. The gate value *g*_*v*_ ∈ (0, 1) is node-specific: nodes in dense, well-connected regions may rely more on the HGT branch (heterogeneous context), while peripheral or under-annotated nodes may rely more on the PNA branch (degree-aware topology).

The GNN is trained jointly on a link prediction objective and an embedding reconstruction objective:

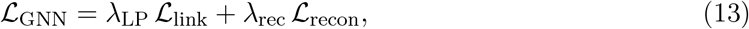

where ℒ _link_ is the binary cross-entropy link prediction loss over sampled positive/negative edges (analogous to Eq. 5), and ℒ _recon_ is a mean squared error reconstruction loss that anchors the GNN output to the pre-trained ComplEx embeddings:

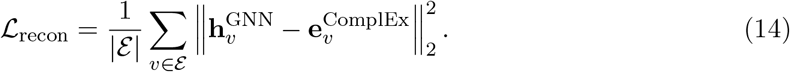

The reconstruction term serves as a regulariser: it prevents the GNN from drifting arbitrarily far from the pre-trained relational structure learned by ComplEx, while still allowing structural neighbourhood information to refine the embeddings. The weighting coefficients *λ*_LP_ and *λ*_rec_ are set to 1.0 and 0.5 respectively, prioritising link prediction while maintaining embedding stability.

### Five-Dimensional Disease Feature Encoding

For each disease node *d*, five categories of external features are assembled from independent data sources and concatenated into a unified feature vector **f**_*d*_ ∈ ℝ^*F*^ :

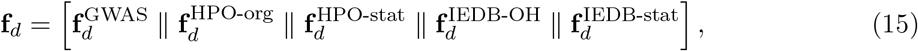

where ∥ denotes concatenation. The five components encode: (1) SNP-level genetic association signals from GWAS; (2) HPO organ-system involvement patterns; (3) HPO phenotype co-occurrence statistics; (4) IEDB antigen one-hot encodings; and (5) IEDB epitope statistical features. Each component is independently standardised with *z*-score normalisation prior to concatenation. All features are keyed by MONDO ontology identifier.

### Three-Source Feature Fusion Network

The core challenge of the fusion stage is that KG embeddings, GNN embeddings, and the hand-crafted disease feature vector **f**_*d*_ live in different representational spaces and carry complementary, non-redundant biological information. A simple concatenation would allow the model to ignore any one source; instead, we employ a **MultiSourceFusionNetwork** with a tri-source attention gate that dynamically weights each source for each individual disease node.

Given the KG embedding **e**_*d*_ ∈ ℝ^*d*^, the GNN embedding **g**_*d*_ ∈ ℝ^*d*^, and the disease feature vector **f**_*d*_ ∈ ℝ^*F*^ for disease *d*, the network first projects each source into a common *d*-dimensional space:

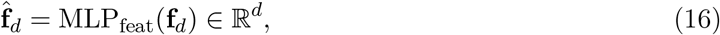

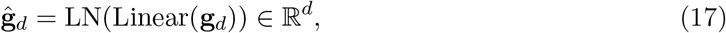

where MLP_feat_ is a multi-layer perceptron with BatchNorm and dropout applied to the heterogeneous feature vector (which mixes binary, continuous, and statistical sub-modules), and LN denotes Layer Normalisation applied to the linear projection of the GNN embedding. The KG embedding **e**_*d*_ is used directly without further projection, as it already resides in the target embedding space.

The three projected representations are then concatenated and passed through a **tri-source gate** that produces source-specific attention weights:

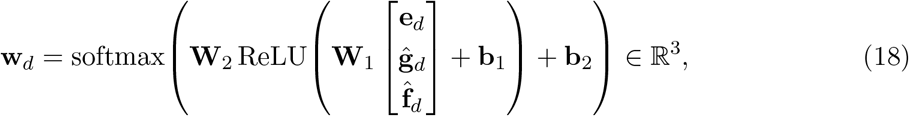

where 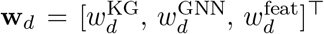 with 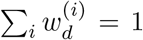. The softmax normalisation ensures that the three weights compete as a probability simplex, allowing the network to identify which information source is most informative for each disease. For example, a disease with rich GWAS and IEDB data but sparse graph connectivity would receive a higher 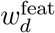 weight. The fused representation is:

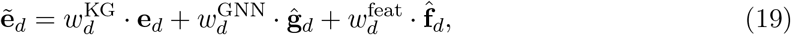

followed by Layer Normalisation to stabilise the scale of the output. The fusion network is trained to minimise the mean squared error between the fused disease embedding and the target pre-trained KG embedding, ensuring that fusion enhances rather than distorts the learned relational structure:

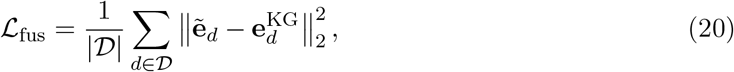

where 𝒟is the set of disease nodes with available external features.

When GNN embeddings are unavailable (e.g., when PyTorch Geometric is not installed), the model degrades gracefully to a **two-source FeatureFusionNetwork** using a single element-wise gating operation:

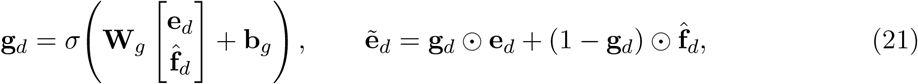

where ⊙ denotes element-wise multiplication. In this fallback mode, **g**_*d*_ ∈ (0, 1)^*d*^ acts as a per-dimension soft switch between the KG embedding and the projected feature vector.

### Target Scoring

The final scoring pipeline combines three complementary signals to rank all candidate gene/protein targets against a query disease group. For each candidate target *t* ∈ *𝒯*_cand_ and disease group *D*_*q*_ (e.g., the set of IBD disease nodes comprising ulcerative colitis, Crohn’s disease, and related entities), three scoring components are computed as follows.

#### KG score

The ComplEx scoring function (Eq. 4) is evaluated for all (*d, r, t*) triples across all disease nodes *d* ∈ *D*_*q*_ and biologically relevant relation types *r* ∈ *ℛ*_bio_. The maximum score is taken to capture the strongest evidence of association, regardless of which specific relation type or disease subtype provides it:

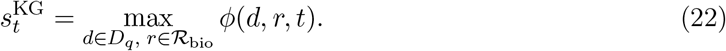

#### Fusion score

Cosine similarity is computed between the fused disease embedding 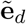 (which integrates KG, GNN, and feature information) and the KG embedding of the target, averaged across the disease group to produce a robust consensus signal:

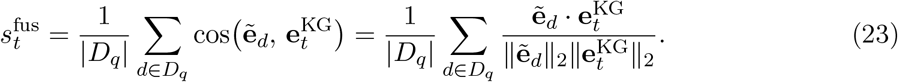

#### GNN score

Cosine similarity is recomputed using the GNN embeddings directly, capturing the structural neighbourhood context independently of the fusion step:

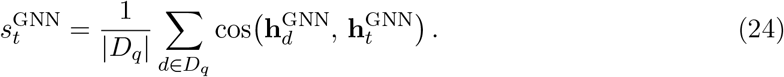

#### Combined score

The three components are aggregated with fixed weights that reflect our relative confidence in each signal source. When all three are available, the KG score receives the highest weight (0.4) as the most directly interpretable biological signal, while GNN and fusion scores each receive 0.3:

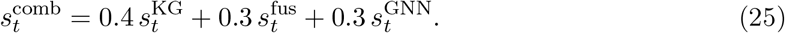

Prior to combination, KG scores are min-max normalised across all candidate targets to align their range with the cosine similarity-based fusion and GNN scores, which naturally lie in [−1, 1].

### Novelty-Penalised Scoring

A known limitation of KG-based scoring is that well-studied, high-degree nodes accumulate spuriously high scores by virtue of their dense connectivity rather than true biological relevance. In immuneKG, prominent hub genes such as TNF or IL6 are connected to thousands of triples, giving them an intrinsic scoring advantage that would systematically suppress dark-target candidates. To counteract this bias and shift the discovery orientation from well-validated hotspots toward underexplored candidates, we apply a degree-based novelty penalty to the combined score:

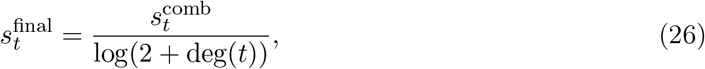

where deg(*t*) is the total degree of target node *t* (sum of in-degree and out-degree in the full KG). The logarithmic denominator provides a smooth, sublinear penalty: a gene with degree 1,000 is penalised by log(1,002) ≈ 6.9, while one with degree 10 is penalised by only log(12) ≈ 2.5. The base term of 2 ensures a finite, non-zero denominator even for isolated nodes with deg(*t*) = 0.

To quantitatively assess the novelty composition of a ranked candidate list, we define the **Average Popularity** (AP) metric at rank *K*:

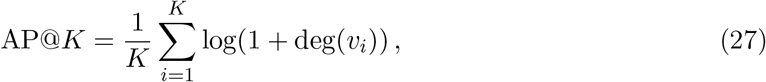

where *v*_*i*_ is the *i*-th ranked target. AP@*K* measures the average graph connectivity (a proxy for how well-studied the targets are) of the top-*K* predictions: a lower AP@*K* value indicates a list enriched in less well-connected, more novel candidates. This metric allows direct comparison of the novelty balance between different scoring strategies (e.g., with and without the degree-based penalty).

### Training Protocol

The three training stages are performed sequentially. In Stage 1, the ComplEx model is trained for 300 epochs with the Adam optimiser (learning rate 10^−3^, batch size 512) using the pre-partitioned train/valid/test split. In Stage 2, the HeteroPNA-Attn GNN is trained for 200 epochs with the combined objective (Eq. 13), initialised from the frozen ComplEx embeddings (*λ*_LP_ = 1.0, *λ*_rec_ = 0.5). In Stage 3, the MultiSourceFusionNetwork is trained for 100 epochs with the MSE objective (Eq. 20), with early stopping triggered after 10 epochs of non-decreasing validation loss. All experiments use random seed 2026 to ensure reproducibility without seed engineering. Models are implemented in PyTorch; KG training uses the PyKEEN library [81]; GNN components are built with PyTorch Geometric [82].

## Discussion

Precise definition of relation types is central to constructing biomedical knowledge graphs with genuine utility for downstream target discovery. Many existing graphs rely on coarse-grained semantic descriptors such as “associated_with,” which lack the mechanistic specificity required for causal reasoning in precision drug development [18, 41]. immuneKG addresses this by enforcing deep semantic granularity in all four novel relation types, each carrying an explicit directional biological interpretation.

Despite this progress, two notable data sparsity challenges remain. First, the `DrIc` (drug–immune cell) relation type suffers from limited coverage: while molecular-level pharmacological data are abundant, systematic records of drug-induced modulation of immune microenvironments at the cellular level remain scarce in current databases [46]. Second, the `IcIm` (immune cell–disease) relation requires richer curation: existing databases predominantly encode direct protein–disease associations while underrepresenting immune cells as pathological intermediaries [47]. These cross-scale data gaps—from molecular to cellular to histopathological levels—limit the graph’s expressive capacity in complex immune-mediated diseases. Future integration of single-cell sequencing data from the Human Cell Atlas [90] and disease-specific single-cell studies in IMIDs [94, 95], together with dedicated immunological interaction databases, will be critical for increasing the density of these two relation types and thereby improving novelty score performance.

We further observed that even within immune-related diseases, certain features remain disease-specific. Because the KG explicitly encodes (Disease)–[ASSOCIATED_WITH]–(Gene) relationships via relational edges, it is unnecessary to redundantly enumerate thousands of core genes within disease node feature vectors. Notably, disease-stratified differentially expressed genes (DEGs) are not incorporated into the feature representation—a key structural advantage of KG-based learning frameworks, as it effectively avoids the dimensional explosion inherent to traditional one-hot encoding schemes. In GNNs, the disease-specific node features complement the graph’s relational edges in a synergistic manner, enabling richer biological context when predicting novel therapeutic targets.

Feature extraction for the autoimmune antibody spectrum presented a methodological choice between two strategies: a clinically driven approach based on gold-standard autoantibody profiles, versus a data-driven approach leveraging antigen frequency statistics from the IEDB database to extract the top-45 most frequent antigens. Given that our downstream objectives prioritise dark-target discovery rather than reinforcement of existing hotspots, the data-driven strategy was selected for its broader antigenic landscape coverage.

### Concept of disease targets versus drug targets

Throughout this study, the term *target* refers specifically to disease targets, distinct from the more commonly cited drug targets. With advances in AI-driven molecular design, traditionally undruggable targets are progressively becoming tractable. For example, the PEGASUS AI model learns design rules for cell-permeable macrocyclic peptides, enabling effective engagement of intracellular targets that were historically inaccessible [91]. Similarly, CANDID-CNS predicts blood–brain barrier (BBB) permeability of small molecules in beyond-Rule-of-Five (bRo5) chemical space, expanding the feasibility of oral neurotherapeutics [92]. A disease-target–centric perspective therefore offers a more forward-looking conceptual framework, shifting emphasis from classical druggability constraints toward biological relevance and disease-network positioning.

### Cold-start entity analysis

In evaluating model performance, cold-start entities—those absent from the training set but appearing in validation or test triples—represent a key challenge for knowledge graph completion. We identified 10 unique cold-start entities following dataset partitioning using random seed 2026 (no seed engineering was applied). Their distribution is detailed in Table 12.

**Table 12:**
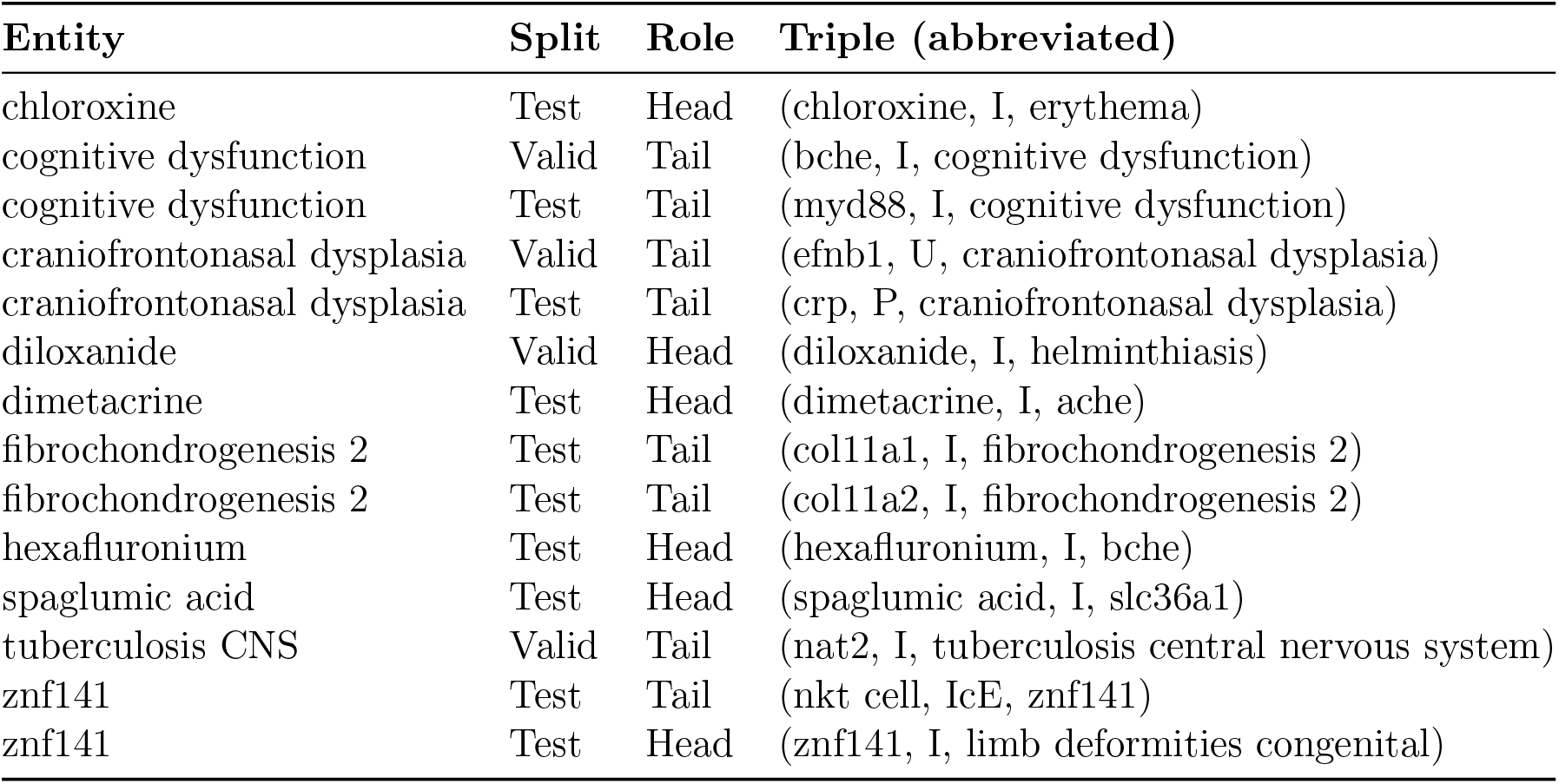
Cold-start entities identified in the immuneKG validation and test sets. “Head” and “Tail” indicate whether the entity appears as the head or tail entity in the respective triple. Entities appearing in both sets are indicated.

Although these entities appear infrequently (typically 1–2 times each) in the evaluation sets, the absence of pre-trained embedding vectors for them disproportionately impacts Hits@1 and MRR metrics. Their presence underscores the importance of multimodal feature-based initialisation (e.g., SMILES or protein sequence embeddings) for handling novel entities in specialised biomedical KGs such as immuneKG.

### On post-hoc rationalisation in AI-driven drug discovery

Although AI-driven drug discovery holds transformative potential, the field faces a systemic challenge: the widespread presence of post-hoc rationalisation. Some studies retrospectively adjust model parameters to fit known experimental outcomes, constructing logically coherent narratives while obscuring generalisation deficiencies and risking metric inflation through implicit data leakage. A further concern is selective experimental validation—reporting only biologically meaningful hits among the top 30–50 scored targets—which constitutes a survivorship-biased paradigm that systematically over-estimates model efficacy. In contrast, this study integrates biologically grounded immune-disease feature embeddings with a highly adaptive deep learning architecture and adopts a strict model-first validation strategy: optimising Hits@1 as the core metric better approximates industrial target discovery needs and structurally eliminates selection bias, representing genuine model-guided target discovery in which biological insight emerges from model inference rather than retrospective narrative construction.

## Abbreviations

SNP: Single Nucleotide Polymorphism
GWAS: Genome-Wide Association Studies
IBD: Inflammatory Bowel Disease
EFO: Experimental Factor Ontology
Chr: Chromosome
LD: Linkage Disequilibrium
BKG: Biomedical Knowledge Graph
OMIM: Online Mendelian Inheritance in Man
Mondo: Mondo Disease Ontology
HLA: Human Leukocyte Antigen
ADs: Autoimmune Diseases
OR: Odds Ratio
Hpoa: Human Phenotype Annotation
CD: Crohn’s Disease
UC: Ulcerative Colitis
HRGAT: Heterogeneous Graph Attention Network
UMAP: Uniform Manifold Approximation and Projection
IMIDs: Immune-Mediated Inflammatory Diseases
KGE: Knowledge Graph Embedding
GNN: Graph Neural Network
PNA: Principal Neighbourhood Aggregation
HGT: Heterogeneous Graph Transformer
TPMI: Taiwan Precision Medicine Initiative.

## Data Availability

The raw results files for the IBD case study and the pre-trained model weights are deposited in a Google Drive repository at the following link: results — Google Drive. The Clarivate Cortellis Phase II+ clinical enrichment reference table used for validation is available as a Google Sheets document: Feb_28_2026_inflammatory_bowel_disease — Google Sheets. The figure generation scripts used to produce all visualisations in this manuscript are provided as a runnable Google Colab notebook: immuneKG — Google Colab. Users should upload the local results folder and link it via the provided Drive path to reproduce all figures. For additional requests or further assistance, please contact the corresponding author at.

## Code Availability

The full source code of the immuneKG project, including data preprocessing pipelines, graph construction scripts, model training routines, and the immune cell structural contribution analy-sis module (`explain_immunecell.py`), is publicly available on GitHub at https://github.com/YaowenYe/immuneKG.

## Acknowledgement

The authors express their deepest gratitude to Professor Dengming Ming for his steadfast support and invaluable guidance throughout this research. Beyond his significant contributions to refining the linguistic quality of this manuscript, his rigorous academic standards and mentorship have been instrumental in shaping the conceptual framework of this work.

With equal sincerity, Yaowen Ye further extends a profound and personal acknowledgment to Ms. Jia Chen of Jingjiang No. 1 Senior High School. The bond between Y.Y. and Ms. Chen was forged at a senior high school in China, where she served as a beloved biology teacher. During this formative period, Y.Y. possessed an insatiable curiosity and a persistent desire for knowledge in life sciences. In response to Y.Y.’s various unconventional biological inquiries, Ms. Chen conducted extensive research to provide rigorous, evidence-based answers that reflected the true complexities of the biological field, never dismissing those questions for being outside the scope of standardized curricula—a visualization to her profound respect for a student’s vigorous intellectual exploration. Despite the limitations of the regional examination scoring scheme at the special time (Jiangsu, China, 2008-2019), which did not fully incorporate Biology into the competitive credit system, Ms. Chen’s unwavering encouragement of such curiosity has become a perpetual beacon. This early enlightenment, now amplified by the professional mentorship of Professor Ming, continues to guide Y.Y. on his path of persistent exploration within life sciences.

